# Automated culture and monitoring of a high-throughput human heart-on-a-chip

**DOI:** 10.64898/2026.03.11.711145

**Authors:** Bryan G. Schellberg, Nolan T. Burson, Joshua Gomes, Guohao Dai, Abigail N. Koppes, Ryan A. Koppes

## Abstract

Organ chips offer a disruptive innovation to study human diseases with tissue-specific resolution within a predictable and tunable in vitro environment. However, these platform technologies have for the most part failed to translate to broad use in the private sector due to a lack of high-throughput, user-friendly platforms. Here we present an automated high-throughput organ chip seeded with iPSC-derived cardiomyocytes transduced with GCaMP6f and interface with translational technologies to bridge the current academia-industry gap. Cardiomyocytes were seeded on-chip fully hands-free using an entry-level fluid handling robot to significantly reduce user handling requirements. Pipette interfaces were paramount to facilitating seeding and feeding through improved tolerances for establishing a functional connection to dispense and collect small fluidic volumes. Following successful seeding, GCaMP6f activity on-chip was monitored with our automated, non-invasive fiber-optic sensing platform. We show a significant decrease in cardiomyocyte beat rate in response to decreased ambient culture temperature using data collected with our optical sensing platform. This study provides a potential translational blueprint for academia-industry partnership toward broad adoption of organ chip technology in drug development and disease modeling.

## Main

Over the last 15 years, microphysiological systems (MPS) or organ chips have emerged as a promising culturing system compared to traditional 2D *in vitro* models. Organ chips enable the use of increasingly sophisticated approaches for disease modeling and drug development including 3D cell culture, complex cell-cell and cell-matrix interactions, and biophysical stimuli such as fluidic shear stress or tunable matrix stiffness^1-4^. These devices have been used to recapitulate numerous organ-specific functions *in vitro*, including the gut epithelial barrier, blood brain barrier, and liver-kidney system^5-7^. For spatial control over cell arrangement, organ chips may be engineered to house cell types in specifically tailored culture environments. Adherent barrier cells can be cultured in 2D on a supporting membrane or extracellular matrix, neurons may be encapsulated in defined 3D chambers, and immune cell migration may be facilitated using interconnecting chambers or channels^8-10^.

The clinical relevance of MPSs has been further improved using human cells instead of animal cells such as recent advancements in human stem cell differentiation. There have been multiple demonstrations of iPSC-derived cell types cultured on-chip, ranging from direct seeding of differentiated cells to complete on-chip differentiation^11-13^. The broadening availability of iPSC-derived cell sources creates unique opportunities for fully humanized organ chip models with unmatched potential for disease modeling, preclinical pharmaceutical screening, and individualized medicine. However, there is a balance between the degree of complexity and ease of use. Culture models that require complex handling, intricate seeding processes, and technical mastery greatly limit the adoption and ability to scale. For pre-translational testing, it is important to ensure that a simplified platform scales well before multiple cell types are introduced. The ultimate goal is to build physiologically accurate platforms that meet the throughput demands of industry^14-17^.

*In vitro* models are only as powerful as their functional outputs and measurements. MPS designs in recent years have been outfitted with external hardware for either improved physiological mimicry or data acquisition^18^. Optical or electrochemical elements have been integrated on-chip for rich data outputs relevant to the physiological model of choice. For example, electrodes have been patterned on-chip for trans-epithelial/endothelial electrical resistance (TEER) measurements in a host of barrier tissue models^19-21^. Patterned microelectrode arrays have also been used to stimulate and sense extracellular voltages of contracting muscle or depolarizing neurons on-chip, highlighting the technical capabilities of current organ-chip technologies^22,23^. However, while integrated electrochemical approaches yield unmatched sensitivity and signal fidelity, electrodes are susceptible to fouling, degradation, and saturation, limiting their application to expensive, single-use, low-throughput sensing approaches^24,25^. The high cost of these chips as well as the data acquisition systems hinders broader adoption.

Despite significant advancement in electrochemical and optical sensor integration into organ chips, there still exists a lack of high-throughput, real-time sensing approaches that meet the needs of industry stakeholders^15,26^. Specifically, pharmaceutical companies have invested significantly into their existing drug screening pipelines and would require that any organ chip device will integrate directly into their established infrastructure^27^. Some commercial organ chip companies, such as Mimetas and CN Bio, have adopted a multi-well plate format for their organ chip designs to be directly compatible with existing analysis tools; however, most organ chips require specialized training and tools, which hampers translation potential. Due to the mismatch between current academic and industry standards for device readiness, most sensor-based organ chip devices remain trapped in academic labs^28^.

High content imaging systems have become a workhorse for the pharmaceutical industry^29^. Optical approaches offer direct observation of the cellular microenvironment without requiring embedded hardware or direct contact with the culture. Brightfield and fluorescence-based assays have become the standard approach to characterize MPSs. Live imaging techniques enables bulk sampling of 2D or 3D culture areas with reliable, reusable components, but typically comes at the cost of reduced temporal sensing capability^17,30^. Imaging equipment is compatible with standard culture ware materials including polystyrene, glass, and PDMS. However, the cost of high content imaging systems has limited their broad accessibility.

Robotic liquid handlers are useful in ensuring reproducible handling and consistent quality; however, most organ chip designs are not amenable to direct integration with these devices. Non-standard chip footprints and narrow fluidic access ports makes interfacing organ chips with automation tools a significant challenge, requiring liquid handlers with micron-level tolerances capable of running specialized protocols^25,31^. Here, we combine a multi-well organ-chip (throughput) seeded with hiPSC-CMs (humanized model), pipette interfaces (automated liquid handling integration), and fiber-optic sensing (automated real-time data) to create an organ-chip platform that meets the needs of both industry and academia.

## Results

### Multi-well organ chip design and interfacing with automation devices

Our previously established layer-by-layer heart-on-a-chip design^2^ was expanded, fitting 12 independent culture areas into a standard well-plate footprint (Figure 1a-c). Standardized dimensions (127 x 85 mm) allow for direct integration with current industrial automation tools and devices, including motorized microscope stages. Additionally, the top layer of our MPS was modified with commercially available pipette interfaces to enable automated on-chip seeding and media exchange via a fluid handling robot. The multi-well MPS design may be readily modified for additional culture models, acting as a blueprint for high-throughput, automation approaches using organ chips^**32**^.

**Fig. 1:**
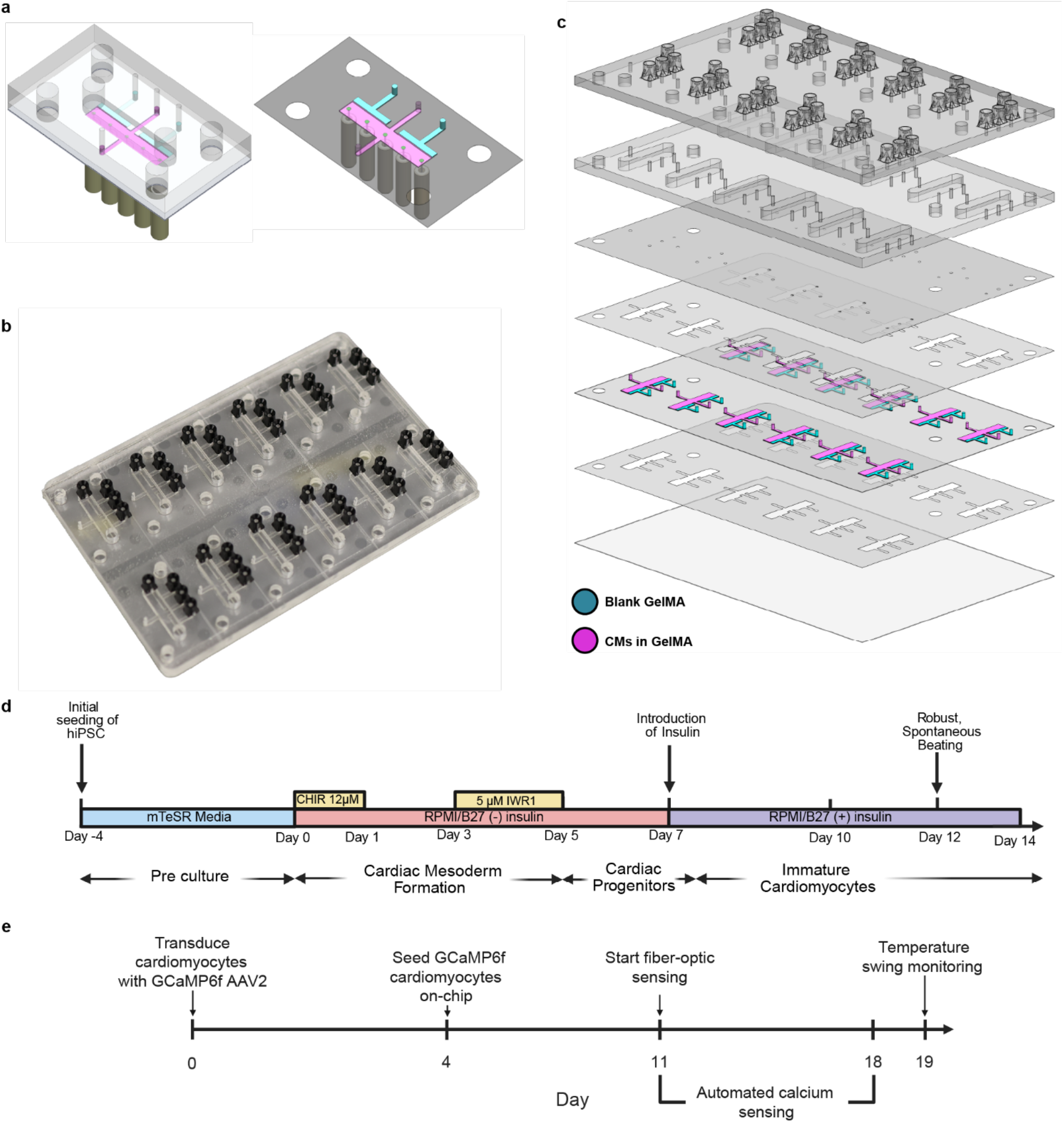
Development of an automated human cardiac MPS. a) 3D rendering of a single MPS well (left) and a cutaway showing the cell culture area within the organ chip (right). The cardiomyocyte culture chamber is shown in magenta, and the blank GelMA is shown in cyan. Fiber optic cannula are positioned underneath the MPS, with approximate illumination volumes illustrated in green. b) Photograph of the multi-well MPS platform. c) An exploded view of the multi-well organ chip, showcasing the individual layers stacked together to assemble the MPS. d) Culture timeline used to differentiate iPSCs to spontaneously contracting cardiomyocytes in 14 days. e) Transduction timeline for stable GCaMP6f expression in iPSC-derived cardiomyocytes. Transduced cardiomyocytes were then seeded in our MPSs and used for fiber-optic sensing.

### hiPSC-derived cardiomyocytes robustly express GCaMP6f in a three-dimensional heart-on-a-chip model

Following an established protocol^33^, we successfully differentiated iPSCs to spontaneously beating cardiomyocytes in 14 days (Figure 1d). After successful differentiation, the cardiomyocytes were transduced with a prepared GCaMP6f adeno-associated virus in a well plate prior to seeding on-chip. The cardiomyocytes robustly express GCaMP6f as early as 24 hours post-transduction, as observed through calcium flux during spontaneous contraction (Movie S1). Following a 7-day recovery period post transduction, the cardiomyocytes were then seeded on-chip, suspended in three-dimensions by a 5% (w/v) gelatin methacrylate (GelMA) hydrogel matrix (Figure 1e). Following a 3-day grow-in period on-chip, spontaneous beating, as well as GCaMP6f expression, was observed on-chip (Movie S2).

### Fluidic pipette interfaces enable integration of organ chip platforms into high-throughput automation workflows

Our engineered organ chip successfully interfaces with a commercial fluid handling robot, allowing for a fully automated cell culture workflow, from 3D cell seeding to daily culture maintenance (Figure 2a). Pipette interfaces were laser welded to the top layer of our MPS, allowing for a reliable temporary connection between the automated liquid handling system pipettor and the fluidic channels within the MPS. The pipette interfaces feature a conical lead-in that accommodated slight misalignment in the X-Y plane, ensuring the pipette was always aligned to the elastomer seal, even when the MPS was not perfectly aligned to the robotic liquid handler (Figure 2b,c). The elastomer seal within the pipette interface also accommodated the common errors of positioning the pipette tip in the Z axis. Combined, the pipette interface enabled the automated liquid handler to drive flow within our MPS, facilitating both cell seeding and daily culture (Figure 2d).

**Fig. 2:**
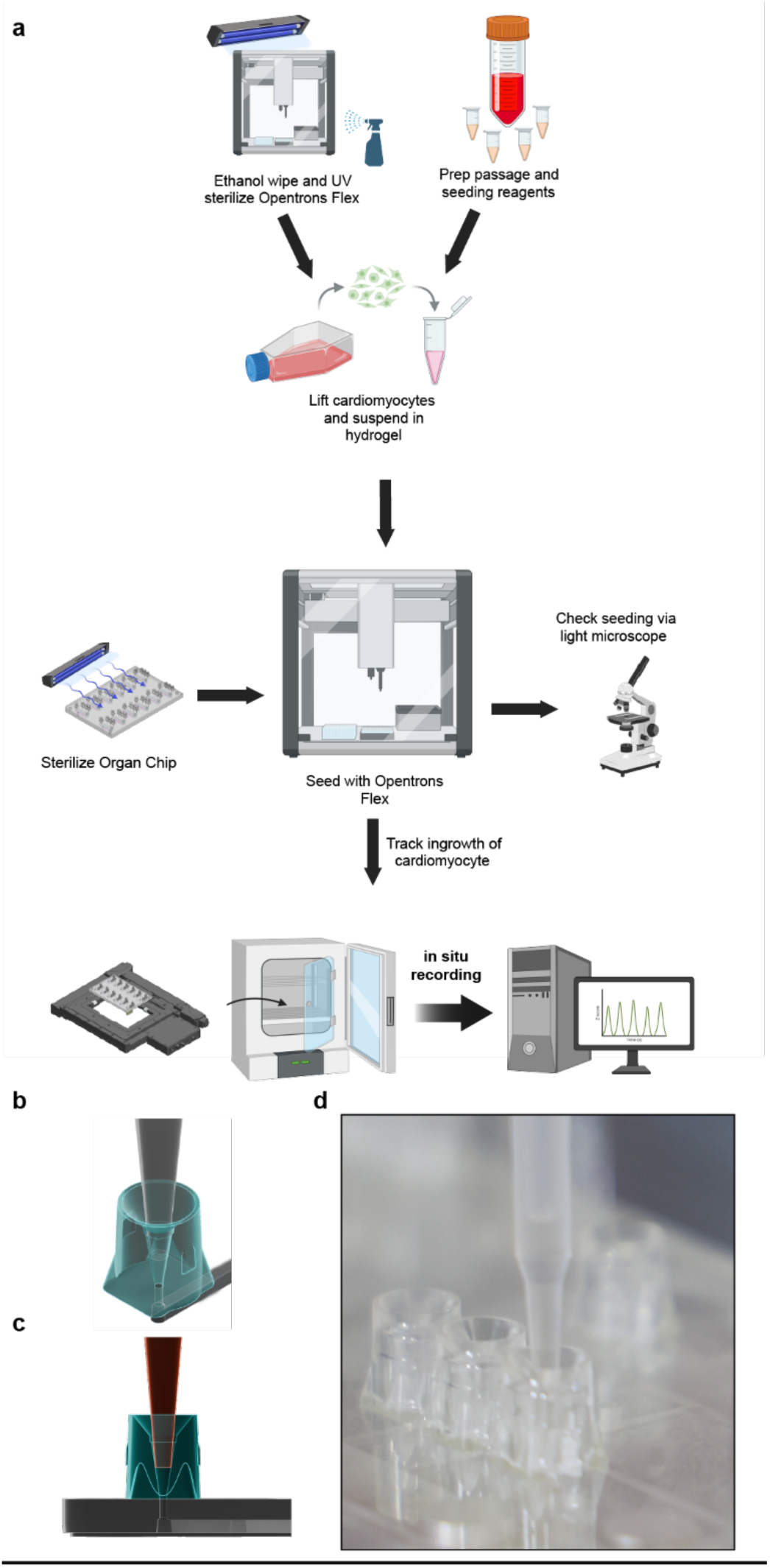
Automated cell seeding and maintenance enabled by a fluidic handling robot and pipette interfaces. a) Workflow used for seeding iPSC-derived cardiomyocytes into our multi-well MPS. MPS preparation and cell work was completed concurrently, as shown, to minimize the total time between cell resuspension in the GelMA hydrogel matrix and seeding on-chip. b) Isometric rendering of the pipette interface, shown in cyan, used for automated cell seeding and maintenance of our MPSs. An example pipette tip, shown in grey, forms a fluidic seal at the interface for streamlined fluidic handling. c) Side view rendering of the pipette interface attached to the top layer of our MPS, shown in grey. d) Photograph of automated cell seeding in our MPS using the pipette interfaces.

Using a custom protocol, we seeded approximately 1 million cardiomyocytes, suspended in 25 μL of GelMA hydrogel, into each of the 12 chip wells. The total seeding time was approximately 17 minutes, including initial cell feeding. There was some variation in seeding quality across wells, such as the formation of air bubbles in the culture well, further optimization of seeding volumes and flow rates may significantly reduce these issues (Supplemental Figure 1). Seeding quality did not impact fiber-based recordings of spontaneous beating. Of note, automated seeding performed better than a trained pipette hand, resulting in a reduced the failure rate at the GelPin (hydrogel overspills) from the main cardiac chamber to the accompanying side chambers. All wells in the multi-well organ chip produced usable data despite 18.8% and 8.3% of wells showing minor and major overspills, respectively. These spill rates, while not perfect, are a significant improvement compared to seeding single-well chips by hand, where 61.5% of our chips exhibit some hydrogel overspill (Supplemental Table 1 and 2).

Organ chip media was replenished daily using an automated fluid handling protocol similar to seeding on-chip. The media replacement protocol was approximately 8 minutes long. Despite taking 2-3 times longer than pipetting by hand, automated fluid handling has the distinct advantage of unmatched repeatability. With fully tunable dispense volumes and rates, all wells receive the same volume of media at a steady, low shear rate. While these details are relatively unimportant for typical *in vitro* cultures, cells cultured in organ chips are susceptible to high shear forces due to the microchannels used for media delivery.

### Fiber-optic scanning and organ chip integration

Innovating off our recently validated fiber-optic sensing platform^34^, a new, unembedded fiber optic array was designed to record GCaMP6f emission signal from 5 spatially independent areas in our multi-well MPS (Figure 3a,b). We have further improved sensitivity to detect 10 kDa fluorescent dextran to a limit of 63 nM, compared to our previously reported limit of 1 μM^34^ (Supplemental Figure 2). Narrow bandpass excitation and emission filters ensured that only GCaMP6f emission was detected, reducing signal contamination by external sources. Further, we recorded GCaMP6f emission concurrently, from an identical cell population, using our optical platform and a fluorescence microscope to confirm comparable data output. We observed nearly identical emission waveforms, validating our optical sensing approach for cardiomyocyte monitoring (Supplemental Figure 3). Compact optical sensing components produced a device with a 15 cm x 25 cm footprint that easily fits on a small portable table outside the incubator (Supplemental Figure 4).

**Fig. 3:**
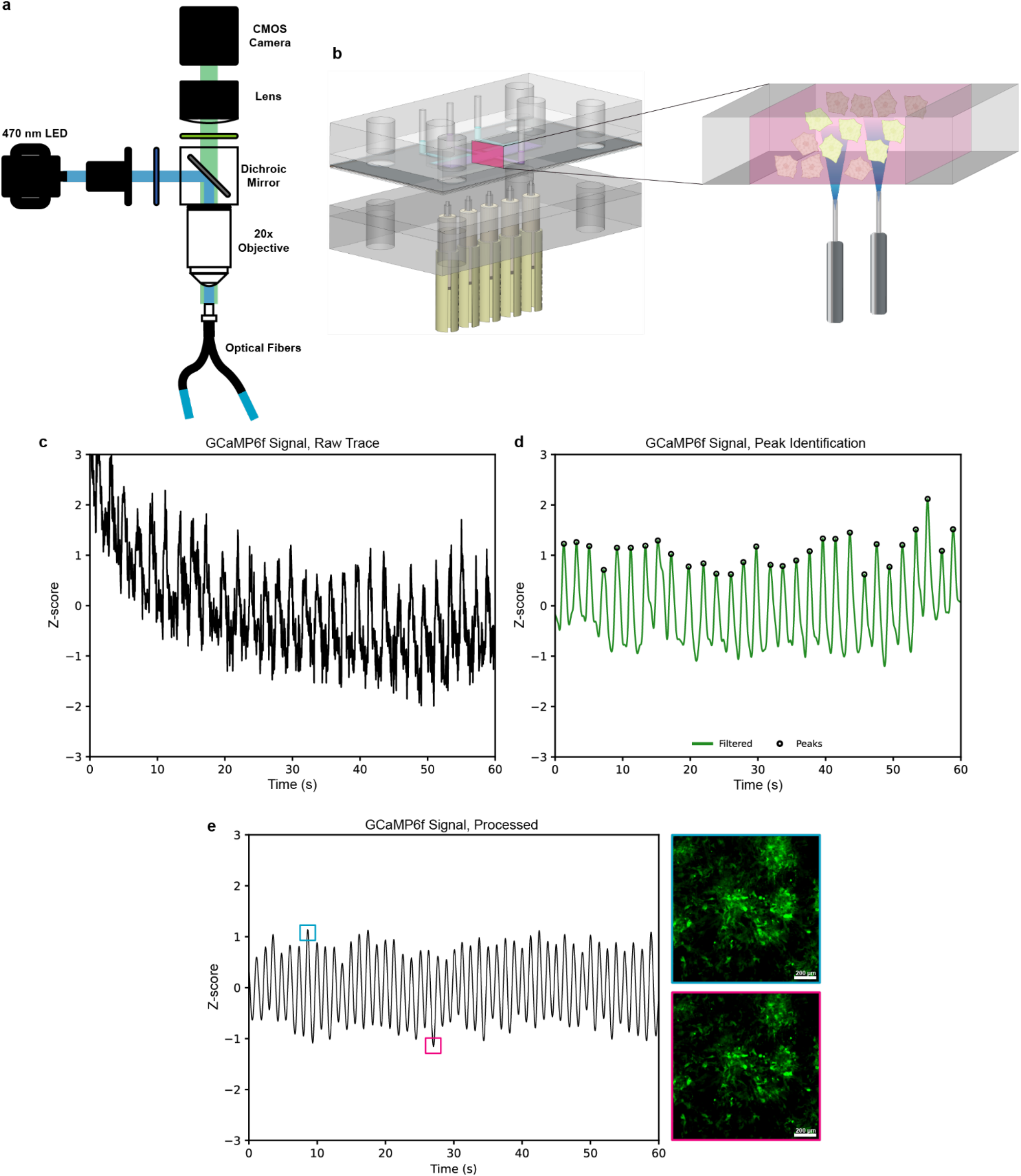
Hands free monitoring of cardiomyocyte beat rate. a) Illustration of the fiber-optic sensing platform used for automated, noninvasive optical sensing from our high-throughput cardiac organ chip. b) A single MPS well (left) and microscopic view (right) of the fiber optic interaction with cardiomyocytes transduced with GCaMP6f and cultured on an MPS. c) Representative Z-scored recording trace collected during in situ recording prior to baseline correction and signal processing. d) Post-processed trace of (c), shown in green, with identified calcium flux peaks corresponding to spontaneous cardiomyocyte contraction shown as black circles. e) Representative processed recording trace, shown in black, with corresponding GCaMP6f expression shown for contracted (blue box, upper right image) and relaxed (magenta box, lower right image) cardiomyocytes on an MPS.

To allow for scanning of all 12 wells using a commercial fiber optic bundle and simplistic hardware design, our MPS was set in a motorized, programmable stage using a standard well-plate adapter. The whole assembly was placed in an incubator for monitoring in a standard culture environment. Fiber-optic cables were routed into the incubator and positioned directly underneath the culture area of interest and held in place by a custom-made jig (Figure 2a). The entire system fits within a single incubator shelf, enabling high-throughput, automated data sensing without sacrificing valuable laboratory space for specialized and potentially bulky sensing equipment. The glass bottom MPS (#2 coverslip) allowed for near-direct (< 1 mm) optical interrogation of the cell culture volume, improving signal-to-noise ratio and reducing possible light scattering, reflection, and dispersion effects on data quality.

### Hands-free, continuous monitoring of cardiomyocyte activity in-situ

To demonstrate the capability of our platform, we recorded whole-MPS GCaMP6f signal across the 12 wells for 7 days without manual manipulation or removal from the CO2 incubator. Our multi-well MPS was set in the motorized stage under normal culture conditions while an Arduino microcontroller and external computer running a custom Python script ran an automated data collection protocol. The sensing platform recorded from 5 spatially independent locations in a single well of our multi-well chip. Data was collected for 60 second epochs. A 20 Hz sampling rate captured highly granular temporal data, as the Nyquist frequency of 10 Hz was significantly higher than the expected (beating/contraction) rate of the cardiomyocytes. Our system is capable of recording the signal from up to a 10 Hz input signal with a 50% duty cycle by recording LED pulses with the fiber optic sensing platform, as expected (Supplemental Figure 5).

Raw fluorescence traces of GCaMP6f were collected using our fiber-optic sensing platform (Figure 3c). Following post-processing and peak detection (Figure 3d,e), we were able to extract a number of signal properties, including beats per minute (BPM), fluorescence rise times, and fluorescence decay times (Figure 4). Due to the size of the dataset collected, comparisons of beating characteristics for both within and across individual wells were evaluated. Further, we used four biological replicates, for a total of 48 MPS wells and analyzed holistic data across all biological and technical replicates (Figure 5).

**Fig. 4:**
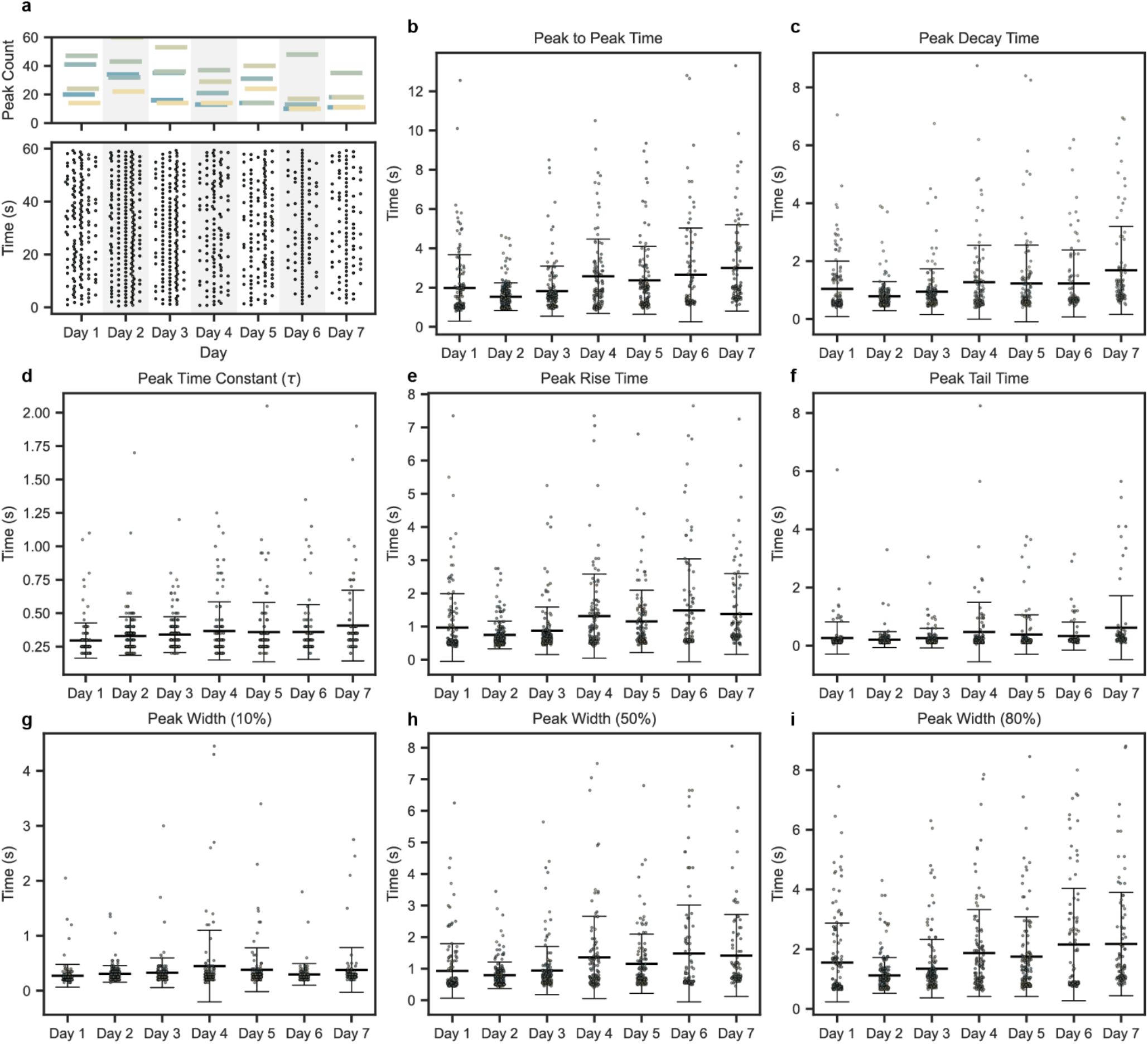
Calcium flux characteristics are measurable via an automated fiber photometry system. a) Representative peak count (top) and spike train (bottom) data collected from a single MPS well across all resting-state recording days. Each bar color represents an individual recording fiber and each data point represents a recorded GCaMP6f spike. Columns of points within each day represent spike trains for individual recording fibers, with Fiber 1 starting on the left to Fiber 5 on the rightmost column. b-i) GCaMP6f peak characteristics extracted from all resting-state recording days. Each data point represents an average value from all 5 fibers in a recording well. Error bars represent mean +/-SD. n=12 wells, m=4 experimental replicates.

**Fig. 5:**
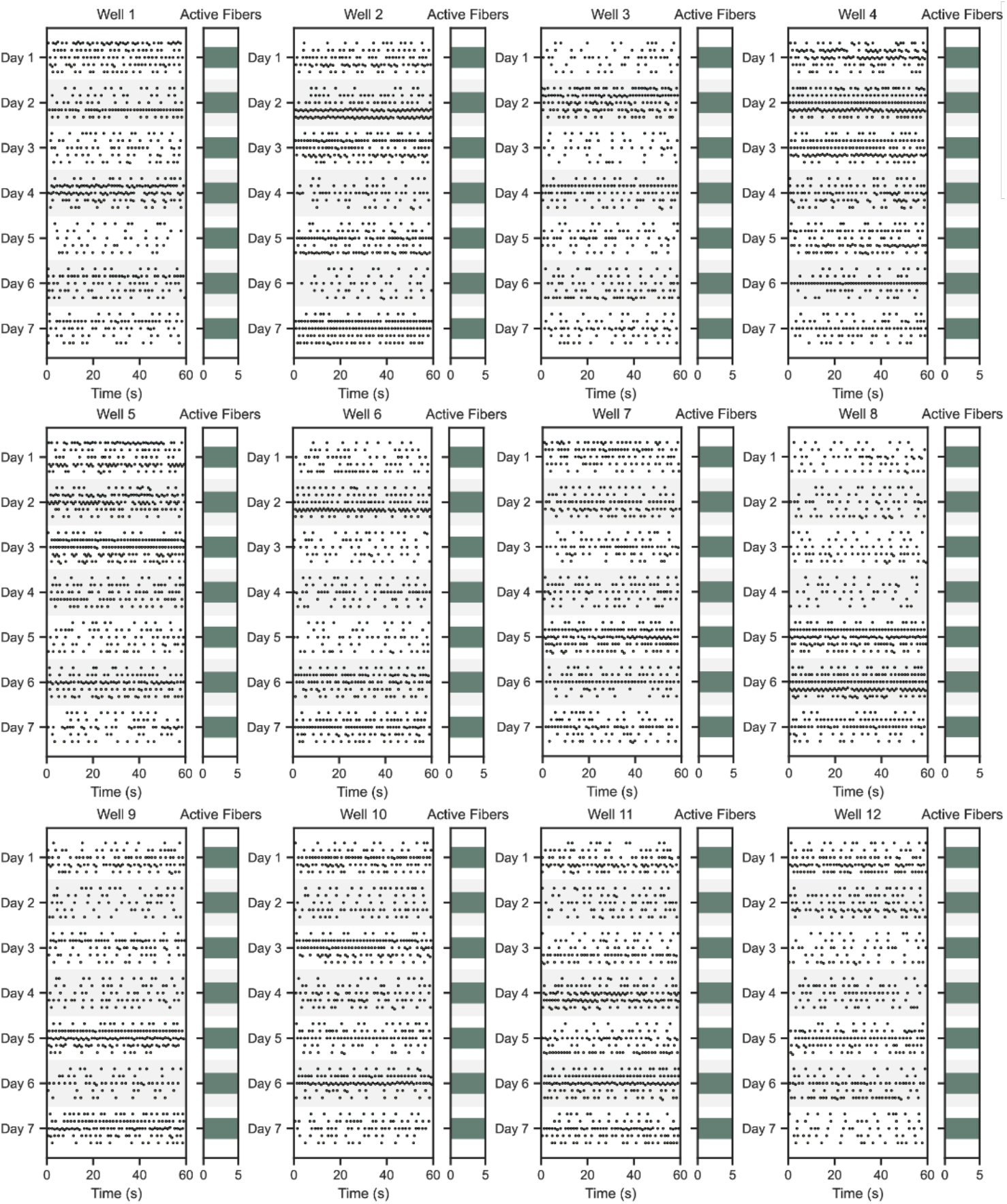
An automated MPS highlights the degree of inherent variability across wells of a single chip with uniform environmental conditions. a) Representative spike trains recorded across all 12 MPS wells during a single resting-state scan. Each data point represents a recorded GCaMP6f spike. Rows of points within each day represent spike trains for individual recording fibers, with Fiber 1 starting on the top to Fiber 5 on the bottom row.

We tracked cardiomyocyte beating over 7 days of culture using our fiber-optic sensing platform, collecting data from each chip once per day. Across all 7 days, we recorded 21.6 ± 2.8 BPM from four organ chips (Figure 6a). There was no significant difference in beat rate across days. Additionally, all fibers were active, with a minimum of 5 BPM, across all organ chips for all recorded days (Figure 6b).

**Fig. 6:**
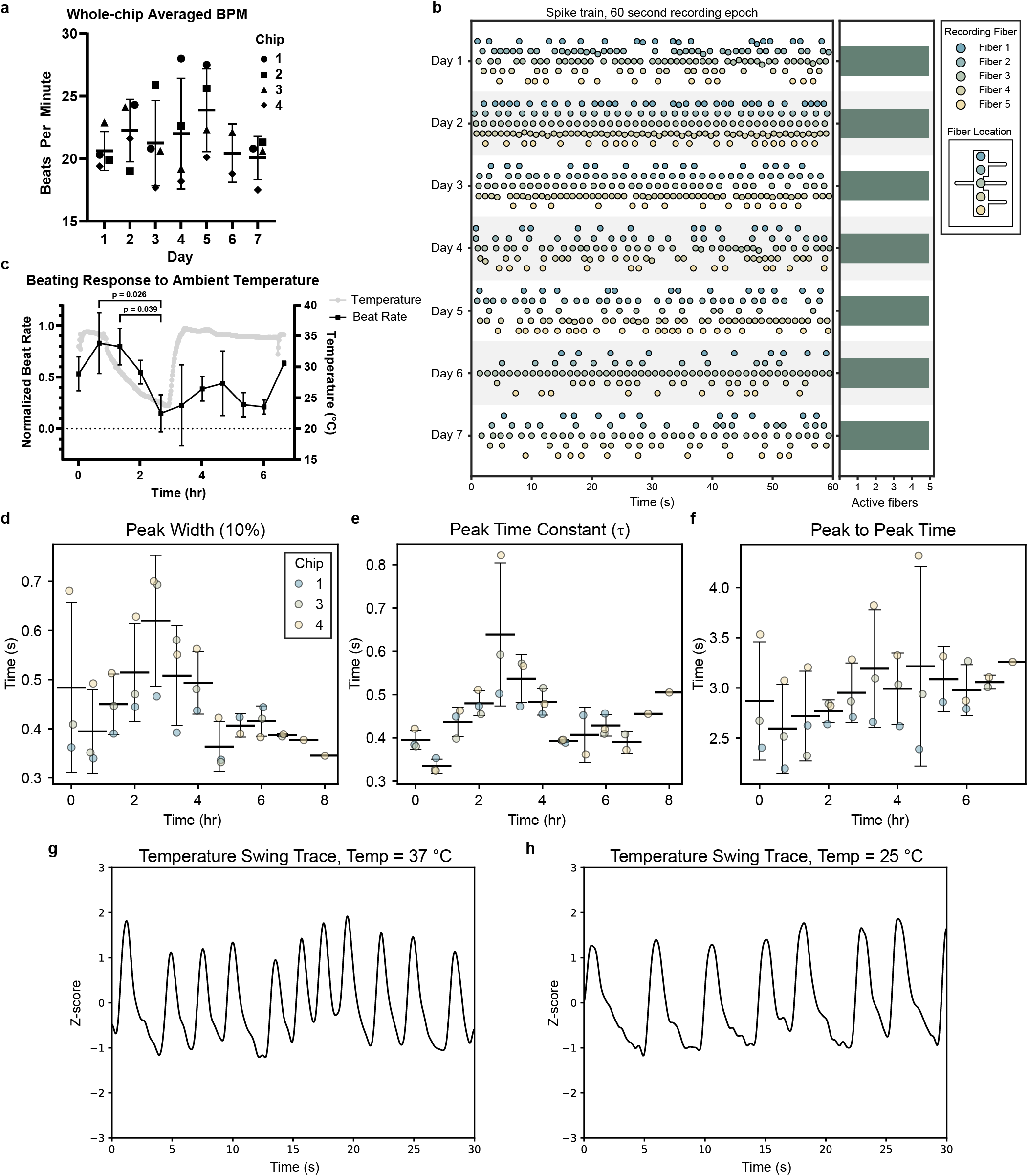
Fiber photometry system captures impact of dynamic temperature changes on beating characteristics. a) Average chip BPM measured across all resting-state recording days. Each data point represents an average BPM across all 12 wells. b) Representative spike train data from a single well across all resting-state recording days. Each data point represents a single GCaMP6f spike. c) Meaured BPM (black) and ambient temperature (grey) as a function of time. BPM data are represented as mean +/-SD and temperature data are represented as the median. n=3 chips. One-way ANOVA with Tukey’s multiple comparisons. d,e,f) 10% peak width, peak time constant values, and peak to peak time of GCaMP6f peaks recorded at each timepoint scan during a temperature swing. Time and temperature values correspond with those shown in (b). g,h) Representative GCaMP6f recording traces from the same well at the first (baseline) and fifth (low temperature) scan.

### Measuring dynamic changes in cardiomyocyte beat rate induced by temperature fluctuations

To demonstrate the temporal monitoring capabilities of our sensing platform, we aimed to track cardiomyocyte beat rate as a function of ambient culture temperature. Over approximately 7 hours, we collected hands-free, beating data from all wells of our multi-well organ chip. The chip was scanned once every 40 minutes, requiring about 10 minutes to complete a full-chip scan. Throughout the recording period, the incubator temperature was modulated to directly impact the observed cardiomyocyte beat rate. Beating rate significantly decreased, compared to early time-points, after the ambient culture temperature reached around 25°C (Figure 6c). This decrease in BPM is expected, as cardiomyocytes are known to be sensitive to temperature fluctuations35,36. Waveform parameters such 10% of peak width and peak time constant (τ) increased as the ambient temperature decreased (Figure 6d,e), while the rise time remained consistent (Supplemental Figure 6a). Additionally, a decrease in variation in the time between peaks as the temperature decreased (Figure 6f). These behaviors trend as expected, as temperature impacts Ca2+ kinetics during relaxation, not the initiation^35^. Although the incubator was able to rapidly restore normal culture temperatures, we did not observe the cardiomyocytes fully rebound from low-temperature exposure. It is likely that extending the recording time, such as monitoring overnight, would show a steady return to pre-experiment beating rates.

## Discussion

Here, we demonstrated the design and implementation of a high-throughput MPS with hands-free sensing and culturing via integration of automation workflows for both seeding and functional measurements. Our design specifications were chosen to address limitations we found when using traditional video microscopy, including limited throughput, the need for expert users, and the analysis of dense datasets. Adoption of a standard well plate footprint allows us to interface with the majority of commercial equipment for fluidic handling and imaging. Expert users are often required for both seeding and imaging, hindering widespread adoption. Further, high content imaging systems and additional support packages add to the financial barrier to the adoption of complex 3D MPSs. As the fiber-optic-based sensing samples a bulk region, we do not require focusing between samples, reducing the workload. Additionally, the use of multiple fibers enables us to collect spatial-temporal data within a single cell chamber, a feature that a traditional microscope cannot achieve. Custom assembled fiber bundles and ferrule connectors hold the potential to dramatically increase this resolution^37^. Towards improving data analysis workflows, fiber-optic-based sensing reduces computational requirements while maintaining quick acquisition times.

The addition of automated cell culture and seeding has been a major improvement in the translatability of this system. Single-well versions saw a ∼60% failure rate in manual seeding by a trained scientist, while the automated workflow has reduced error rates to ∼27%. The high repeatability of automated seeding would allow non-expert users to seed the devices without compromising culture quality. High repeatability of seeding cultures minimizes wasted materials and time, critical when working with valuable cell types. The time required to exchange media and seed the platform was not optimized. The protocol times could be reduced by eliminating extraneous movements and path optimization.

Transduced cardiomyocyte cultures in our MPS declined after 7 days, as evidenced by a decrease in mean beat rate and a decline in culture quality. Previous chip cultures comprised of stem cell-derived cardiomyocytes were repeatably cultured for 30 days (Supplemental Figure 7). Testing the culture limit, a representative MPS exhibited spontaneous contractions for nearly 6 months (Supplemental Figure 8). The transduction of GCaMP6f most likely impacted the metabolic load as well as the viability of the cardiomyocyte cultures implemented for our fiber optic evaluation. While the exact mechanism responsible for the reduction in viability is unknown, we hypothesized that it is a combination of the transition into a three-dimensional culture, the metabolic burden of GCaMP6f, the hydrogel matrix stiffness, and the transduction timeline. Further culture optimizations will be required for long-term experiments, if needed. The current system may be well suited for preliminary dose dependent toxicology data collection.

Current MPS technology faces significant barriers to direct integration into existing drug discovery pipelines, often falling short in ease of use, throughput, reproducibility, and reliability^28^. To fully harness the physiological mimicry of organ chips compared to current *in vitro* and *in vivo* standards, the field must shift toward automation to reduce variability from human error. There is a balance to strike, especially in early design iterations, between biological complexity and scalability when building organ chip systems. To this end, we have developed a simplified, monoculture model that directly integrates into existing, standardized automation tools. We focused on reducing the translational barrier by demonstrating the scalability and automation capabilities of our platform, building out from a single-chip system to a fully automated, 12-well format chip with noninvasive, spatiotemporal data output. Reducing the level of human interaction with organ chip devices will significantly improve device reliability and reproducibility of observed results.

As the pharmaceutical industry increasingly invests in organ chip technology, automation has profound implications. With additional optimization, our cardiomyocyte-based model may act as a preliminary screening tool by facilitating MPS seeding and recordings. By humanizing the drug screening pipeline, there is an opportunity to reduce both false positive and false negative candidates and, as a result, decrease the failure rate of early clinical trials.

## Methods

### Organ-chip fabrication

Organ-chips were assembled layer-by-layer using our previously established laser-cut-and-assembly method ^34,38,39^. Each layer was designed using computer-aided design software (SolidWorks, Adobe Illustrator) and converted to vector format. Individual layers were laser-cut (Epilog Zing) from either transparent, cast polymethyl methacrylate (PMMA, McMaster-Carr, 8560K171) sheets or polyester (PET, McMaster-Carr, 8567K32). PET semi-permeable membranes with 1 μm pore size (It4Ip, ipCellCulture 2000M12/620M103) separated the 3D seeding compartments from the media channels while allowing diffusion of nutrients and cellular waste products across the interface. The uppermost layer containing the fluidic vias was manufactured by milling PMMA sheet stock (Roehm Cyrolite MD-H12) with a CNC machining center (Datron Neo). Pipette interfaces (Parallel Fluidics), were then assembled concentrically above each fluidic via and permanently welded to the uppermost layer using a CNC laser welder (Hymson WS-AT). Individual layers were permanently bonded with pressure-sensitive double-sided tape (966, 3M) as previously described. Large no. 2 cover glasses were (260452, Ted Pella, Inc.) cut to required dimensions with a diamond scribe and used to seal the bottom of the chip and allow for optical access. Organ-chips were assembled by hand, guided by a 3D printed jig to ensure layer alignment. Once assembled, organ-chips were heat pressed at 50°C for 60 seconds under light pressure and placed in a vacuum oven at 50°C for at least 5 days to allow the adhesive tape to sufficiently off-gas and fully cure.

### Fiber-optic sensing setup

Our fiber-optic sensing platform is composed primarily of a CMOS camera, a set of filters and a dichroic mirror to isolate the excitation and emission wavelengths, fiber-optic patch cables, and a 20X objective. The device was engineered to simultaneously deliver excitation light and continuously record emitted signal from 5 spatially arranged locations in each well of our multi-well organ-chip.

The platform was assembled as described below, according to the direction of the beam path. A high-powered 470 nm LED (M470F3, Thorlabs), controlled by a 4-channel LED driver (DC4104, Thorlabs), was coupled via a fiber-optic patch cable (M59L01, Thorlabs) to a collimator (F671SMA-405, Thorlabs). Light shaped by the collimator passed through an excitation filter (470/10 nm, FBH470-10, Thorlabs) held in a quick-release filter holder (SM1-QP, Thorlabs). A dichroic mirror (FL-007031, IDEX) directed the filtered excitation light through the back of a 20X magnification objective (MRH00205, Nikon) and onto the proximal end of a bundled optical fiber (BF7F1LS1, Thorlabs), held in a 5-axis translator (KC1T, Thorlabs). Each distal end of the optical fiber bundle was connected to a fiber-optic cannula (CFMC22L02, Thorlabs) via a ceramic mating sleeve (ADAF1, Thorlabs). The cannula were spatially arranged in a custom laser-cut mount to maintain consistent positioning throughout optical recording. Emitted fluorescence was recovered to the dichroic mirror and directed through an emission filter (520/10 nm, FBH520-10, Thorlabs) held in a quick-release filter holder (SM1-QP, Thorlabs). Filtered fluorescence was focused onto a CMOS camera (BFS-PGE-31S4M-C, FLIR) by an achromatic doublet (AC254-035-A-ML, Thorlabs) and recorded for analysis.

### Automated stage control and image capture

A custom Python script was used to control a motorized, programmable stage (X-ASR100B120B, Zaber) to automate data collection from each multi-well organ-chip using pre-set coordinates. For each recording cycle, the stage randomly traversed the full set of 12 wells, without repeats, pausing at each well for one optical recording epoch. To align our fiber-optic recordings with stage movement, the 4-channel LED driver and CMOS camera were linked to an Arduino microcontroller (Mega 2560 Rev3, Arduino) for trigger control.

At each well, the excitation LED was automatically set to ‘on’, delivering 88 – 131 μW/cm^2^ to each fiber end, exciting GCaMP6f on-chip. Simultaneously, 20Hz square-wave TTL triggers were sent to the CMOS camera to record emitted GCaMP6f fluorescence. Captured images were automatically saved to a pre-named folder on an external laptop connected to the CMOS camera (SpinView v3.2, FLIR).

### Fiber-optic image analysis

Fiber-optic based recordings were analyzed using custom Python scripts to extract relative fluorescence intensity values. The scikit-image package (scikit-image.org; v0.24.0) was used to detect and draw regions of interest (ROI) around each optical-fiber end and extract the mean pixel intensity within each ROI. Calculated values, with timestamps and well identifiers, were exported to a CSV file for additional analysis.

### Peak identification and characterization

Mean intensity values were further processed to allow for identification and characterization of sensed GCaMP6f peaks. Briefly, intensity values for each fiber were subtracted from a fitted logarithmic decay curve to account for LED heat dissipation and photobleaching effects. The fitted curves were then normalized by Z-scoring. The normalized traces were filtered and smoothed using a 1D Gaussian filter with the sigma value set to 3.75. Peaks and corresponding peak characteristics were identified from the filtered data traces using SciPy’s signal processing module (scipy.org; v1.13.1). Processed data, identified peaks, and peak characteristics were exported to a CSV file for data visualization and statistical analysis.

### Cell culture and differentiation

Cardiomyocytes were differentiated from induced pluripotent stem cells (WISCi004-B) following an established protocol using the same stem cell line^33^. To summarize, hiPSCs were cultured to confluency with mTeSR plus media (100-0276, StemCell Technologies) in a 12-well plate (3513, Corning) coated with 1:100 Matrigel GFR (354230, Corning):DMEM/F12 (11320-033, Gibco). Once confluence reached >90%, the cells were switched to differentiation media composed of RPMI1640 (11875093, Gibco) and B27 (-) insulin (A1895601, Gibco) with 12 μM CHIR99021 for 24 hours. Exactly 24 hours after the addition of CHIR99021 (13122, Cayman Chemical, a complete media change was performed and replaced with differentiation media. On day 3, a half media change was performed with fresh differentiation media containing IWR-1 (I0161, Sigma-Aldrich) at 5 μM concentration. On day 5, the media was once again replaced with fresh differentiation media. Day 7 and every 3 days after, the media was changed with RPMI1640 and B27 (17504044, Gibco). Spontaneous contraction began around day 9, and robust beating was reached at day 15, with each well yielding approximately 3 million cells.

### Cardiomyocyte transduction with AAV2-GCaMP6f

Differentiated cardiomyocytes were transduced with prepared adeno-associated virus GCaMP6f (pAAV.CAG.GCaMP6f.WPRE.SV40 (AAV2), <u>addgene.org/100836/</u>, AddGene) to measure Ca^2+^ flux, and consequently, contraction characteristics via our fiber-optic sensing platform. In preparation for cell seeding, a 24-well plate (3526, Corning) was treated with 400 μL of 1:100 Matrigel:DPBS solution overnight. The following day, cardiomyocytes were washed with warm (37°C) DPBS for 5 minutes. After washing, cardiomyocytes were harvested from wells by treatment with 0.25% Trypsin-EDTA (Gibco) for 15-20 minutes at 37°C. After cell detachment, an equal volume of cell culture media (RPMI1640, 10% FBS (16000-069, Gibco), 10 μM ROCK inhibitor (Y0503, Sigma-Aldrich)) was added to each well. Cells were gently triturated to form a single-cell suspension and centrifuged at 200 x g for 5 minutes at room temperature. The supernatant was removed, and the pelleted cells were resuspended in 2 mL of cell culture media; approximately 6×10^6^ cells/mL. Prior to seeding, the Matrigel:DPBS solution was aspirated from each well. Cardiomyocytes were then seeded at a concentration of 8×10^5^ cells per well.

Immediately following seeding, viral particles were suspended in cell culture media and added to each well at a multiplicity of infection of 8.5×10^4^ (6.8×10^10^ genome copies/well) for a final culture volume of 400 μL per well. After 24 hours, the virus-containing media was aspirated and replaced with 1000 μL of fresh cell culture media (RPMI 1640, 10% FBS). Cardiomyocytes were then cultured at 5% CO_2_ and 37°C, replacing media every other day until seeding on-chip.

### Automated organ-chip seeding

In preparation for seeding, multi-well chips were UV sterilized (300 mJ/cm^2^) for 10 minutes on top and bottom sides (Spectrolinker XL-1000, Spectronics Corporation, Westbury NY). Similarly, the deck and pipettes of the fluid handling robot (Opentrons Flex, Opentrons Labworks Inc.) were gently wiped down with 70% ethanol solution and UV sterilized for at least 20 minutes under filtered airflow.

Gelatin methacrylate (GelMA) polymer was synthesized according to previous protocols^40-42^. A 5% w/v GelMA hydrogel solution was prepared by first adding 4 mg of lithium phenyl-2,4,6-trimethylbenzoylphosphinate photoinitiator (LAP, Allevi) to 800 mL of cell culture media containing (RPMI1640, 10% FBS, 10 μM ROCK inhibitor). Once dissolved, 40 mg of GelMA polymer was added to the solution and vigorously shaken to mix. Once fully dissolved, the GelMA solution was wrapped in aluminum foil to protect from light exposure and stored at room temperature until needed for use.

Cardiomyocytes transduced with GCaMP6f were harvested from wells as previously described and centrifuged at 200 x g for 5 minutes. The supernatant was removed, and the pelleted cells were resuspended in 1 mL of cell culture media. The cell suspension was transferred to a 1.5 mL mini centrifuge tube and spun again at 200 x g for 5 minutes. The supernatant was removed, and the cell pellet was carefully resuspended in 340 μL of GelMA solution to achieve an approximate seeding density of 1×10^6^ cells per organ-chip well.

While the cardiomyocytes were prepared for seeding, multi-well chips were treated with oxygen plasma for 90 seconds (Harrick Plasma PDC-001) and moved into position in the fluid handling robot. Using a custom protocol, 7 μL of GelMA solution was automatically seeded into the blank gel ports of the multi-well chip. Once all ports had been seeded, a 10 W UV light (405 nm) box was placed over the chip for 60 seconds to crosslink the GelMA hydrogel. The GelMA seeding was timed such that the gel was fully crosslinked just before the cardiomyocytes-GelMA suspension was prepared.

The cardiomyocyte-GelMA suspension was then immediately placed in position within the fluid handling robot, and the custom cell seeding protocol was started. Each chip well was seeded with approximately 1×10^6^ cardiomyocytes via the appropriate seeding port. The UV light box was again placed over the multi-well chip for 60 seconds to crosslink the GelMA in place. After seeding the blank ports and cardiac chamber, the seeding efficiency was determined via a light microscope, with overspills and airgaps recorded on a paper template of the chip.

Cell culture media was then automatically added by the fluid handling robot via cell feeding pipette interface ports. After adding the culture media, the multi-well chip was removed from the fluid handling robot and cultured at 5% CO_2_ and 37°C for 2 hours. After 2 hours, the culture media was replaced with fresh media, following the automated cell feeding protocol outlined below, to remove trace unreacted photoinitiator and un-crosslinked GelMA.

### Automated organ-chip media replacement

Prior to running the automated media replacement, the fluid handling robot’s deck and pipettors were gently wiped down with 70% ethanol solution and UV sterilized for at least 20 minutes under filtered airflow. Pre-heated aliquots of the appropriate cell culture media were then positioned inside the robot. The multi-well organ-chip was then placed inside the robot, and the custom media replacement protocol was started.

After protocol completion, the chip was removed and cultured at 5% CO_2_ and 37°C.

### Resting-state cardiomyocyte monitoring

Baseline GCaMP6f activity was recorded from cardiomyocytes seeded on-chip using our fiber-optic sensing platform. Base media (RPMI1640, 10% FBS) was replenished 2 hours before each recording to ensure consistent nutrient availability across timepoints. Additionally, the excitation LED was set to maximum power at least 2 hours before recording to photobleach the fiber-optic cables and maintain minimum autofluorescence during recording. The multi-well organ-chip was then placed on the incubated (5% CO_2_, 37°C) programmable stage and our custom data collection script was run to automatically collect resting-state data from all 12 organ-chip wells. Briefly, the programmable stage translated to a randomly selected well position, where the fiber optic platform was triggered to start recording. After recording emitted GCaMP6f fluorescence for 60 seconds at 20 HZ, data collection was paused until the stage moved to the next random well. Recordings were collected from all 12 wells in random order, without repeats, to remove any potential procedural or equipment effects on data quality. Once recordings were completed, the chip was removed from the programmable stage and cultured at 5% CO_2_ and 37°C.

### Temperature swing cardiomyocyte monitoring

To monitor spatiotemporal effects of temperature on measured cardiomyocyte activity, ambient culture temperature was gradually decreased to room temperature (∼25°C), held for an hour at room temperature, and then restored to 37°C. A type K thermocouple was set up to record the temperature within the incubator throughout the recording period, sampling the ambient air temperature every minute. The recording system was prepared in the same manner as the resting-state recordings; however, the full-chip recording scan was scheduled to repeat every 30 minutes to capture temporal changes in cardiomyocyte activity during the temperature swing. Additionally, recording epochs were reduced from 60 seconds to 30 seconds per well to more closely capture whole-chip activity around a recording timepoint. At the end of a recording scan the motorized stage translated to the home position and waited until the next timepoint.

To decrease the ambient culture temperature, the incubator door was propped slightly ajar until equilibrium with the room temperature was reached. After an hour at room temperature, the incubator door was closed, and the ambient culture temperature was allowed to increase by incubator control. Once the temperature stabilized to 37°C, the recordings were continued for an additional 3 hours.

## Data Availability

The data supporting the results of this study and available within the paper and its Supplementary Information. Image files are available upon request from the corresponding author.

## Code Availability

Python code for data collection, data analysis, and Opentrons control can be found at https://github.com/LNNR-and-ABNEL/automated-fiber-optic-sensing-of-GCaMP6f-on-chip with DOI: 10.5281/zenodo.18685793.

## Acknowledgements

This work was supported by the NIGMS MIRA R35 and NASA 21-3DTMPS_2-0037. We would like to thank Dr. Douglas Kim for graciously supplying the prepared adeno-associated virus used in this study.

## Author contributions

**B.G.S./N.T.B**.: Conceptualization, Formal Analysis, Investigation, Methodology, Writing – original draft, Writing – review & editing **B.G.S./N.T.B**.: Conceptualization, Formal Analysis, Investigation, Methodology, Writing – original draft, Writing – review & editing **J.G**.: Conceptualization, Writing – original draft, Writing – review & editing

**G.D**.: Conceptualization, Funding acquisition, Investigation, Writing – review & editing

**A.N.K**.: Conceptualization, Funding acquisition, Investigation, Methodology, Resources, Supervision, Writing – review & editing **R.A.K**.: Conceptualization, Funding acquisition, Investigation, Methodology, Resources, Supervision, Writing – review & editing

