## Supplemental Information for "Automated culture and monitoring of a high-throughput human heart-on-a-chip"

### Automated culture and monitoring of a high-throughput human heart-on-a-chip through academic-industrial partnership: bridging the gap

Bryan G. Schellberg[1*], Nolan T. Burson [1*], Josh Gomes[4], Guohao Dai[2], Abigail N. Koppes [1,2,3] and Ryan A. Koppes [1,2]

**Supplemental Methods:**

**Single Chip seeding and culturing**

Single chips were UV sterilized (300 mJ/cm$^2$) for 10 minutes on the top and bottom sides. While UV sterilizing, a 7.5% w/v GelMA hydrogel solution was prepared by adding 2 mg of LAP to 400 ul of cell culture media containing RPMI160, 10%FBS, and 10μM ROCK inhibitor. Once well mixed, 30 mg of GelMA polymer was added, and the mixture was shaken vigorously while wrapped in aluminum foil. The desired number of cardiomyocytes was lifted and pelleted as previously described.

Pelleted cardiomyocytes were suspended in 80 μL of hydrogel precursor solution to achieve an approximate seeding density of 1x10$^6$ per cardiac chamber. 20 μL of cell suspension was pipetted directly into the center-seeding port. After filling the cardiac chamber, seeded organ chips were placed in a 10W UV light (405nm) box to crosslink for 60 seconds. A light microscope was used to confirm that the hydrogel was completely crosslinked and to check for hydrogel overspills into the side chambers. Empty side ports were filled with blank hydrogel and crosslinked for an additional 60 seconds.

Cell culture media was added to the chip and then replaced 1 hour later to remove unreacted hydrogel components. Daily media exchanges were performed throughout the culturing period. The first 3 days used media containing RPMI1640 and 10% FBS, and every day thereafter used RPMI1640 and B27.

**Immunofluorescent staining**

At the end of the culturing period, culturing media were removed, and a warmed PFA solution (4% in DPBS) was added for 40 minutes. The PFA solution was removed, and DPBS washes were conducted (3 washes, 10 minutes incubation). Cells were perforated with 0.1% Triton X, incubated for 30 minutes, then washed three times with DPBS. Binding sites were blocked using 2.5% goad serum solution for 16-20 hours at 4 °C on a rocker. After blocking, primary antibodies (anti-sarcomeric alpha actinin) were added to a 2.5% goat serum solution and pipetted onto the chip. Primary antibody incubation occurred at 4 °C on a rocker for 16-20 hours. Primary antibodies were washed from the culture using DPBS for 1 hour at 4 °C, on a rocker. Secondary antibody incubation and washes were performed in the same manner as the primary antibodies. After secondary antibody washes, a 1:1000 dilution of DAPI in 2.5% goat serum was added for 10 minutes, followed by three 10-minute wash sets. After the final wash, the chip was ready for imaging or could be stored temporarily with DPBS to ensure the hydrogel does not dry out.

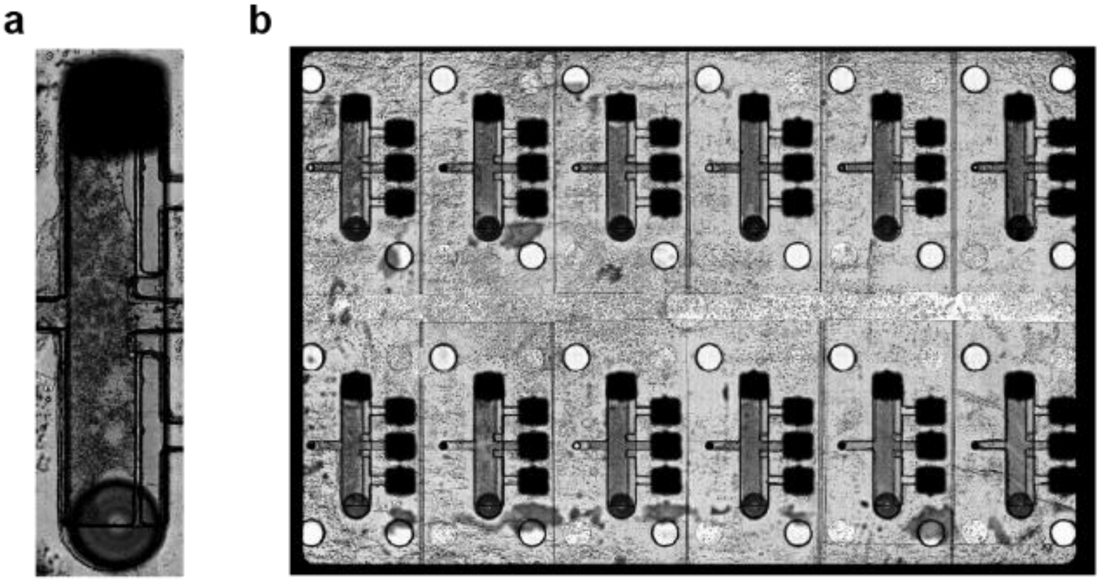


Supplemental Figure 1: a) Tile scan of well A1 showing a small overspill from blank chamber into cardiac chamber b) tile scan of the whole multi-well MPS after 5 days of culturing. Minor seeding errors can be seen such as small overspills from blank chambers into the cardiac chamber and small air pockets at the top of each cardiac chamber. All wells were deemed useable and had little to no impact on data collection.

Supplemental Table 1: Overview of on-chip seeding success using a fully automated seeding protocol conducted with our multi-well organ chips and the Opentrons Flex liquid handling robot. Although we observed both minor and major spillage of GelMA hydrogel from the blank seeding chambers into the cardiomyocyte culture chamber, all wells were still viable for optical sensing. Minor and major leaking were qualitatively assigned to wells where approximately 0-1 $\mu$L or 1-3 $\mu$L of GelMA hydrogel encroached on the cardiomyocyte chamber, respectively.

| Chip Number | Number of seeded wells | Minor GelPin leaking | Major GelPin leaking | Percent usable |
|---|---|---|---|---|
| 1 | 12 | 2 | 0 | 100 |
| 2 | 12 | 3 | 0 | 100 |
| 3 | 12 | 4 | 1 | 100 |
| 4 | 12 | 0 | 3 | 100 |
| Total | 48 | 9 | 4 | |
| % of Wells | 100% | 18.8% | 8.3% | |

Supplemental Table 2: Overview of on-chip seeding success using our previous by-hand seeding protocol conducted with our single-well organ chips. We observed a high failure rate due to GelMA hydrogel from the cardiomyocyte seeding chambers leaking into the neuron culture chamber. Success and failure were assigned whether neurons successfully filled the entire seeding chamber or not. Partial neuron seeding was still considered as a failure, similar to major GelPin leaking described in Table S1.

| Date | Pass GelPIN | Failed GelPIN | Percent success | Percent failure |
|---|---|---|---|---|
| 7/5/2024 | 6 | 3 | 66.7% | 33.3% |
| 8/26/2024 | 2 | 12 | 14.3% | 85.7% |
| 8/29/2024 | 7 | 9 | 43.8% | 56.3% |
| 9/14/2024 | 5 | 8 | 38.5% | 61.5% |
| Total | 20 | 32 | | |
| % Total | 38.5% | 61.5% | | |

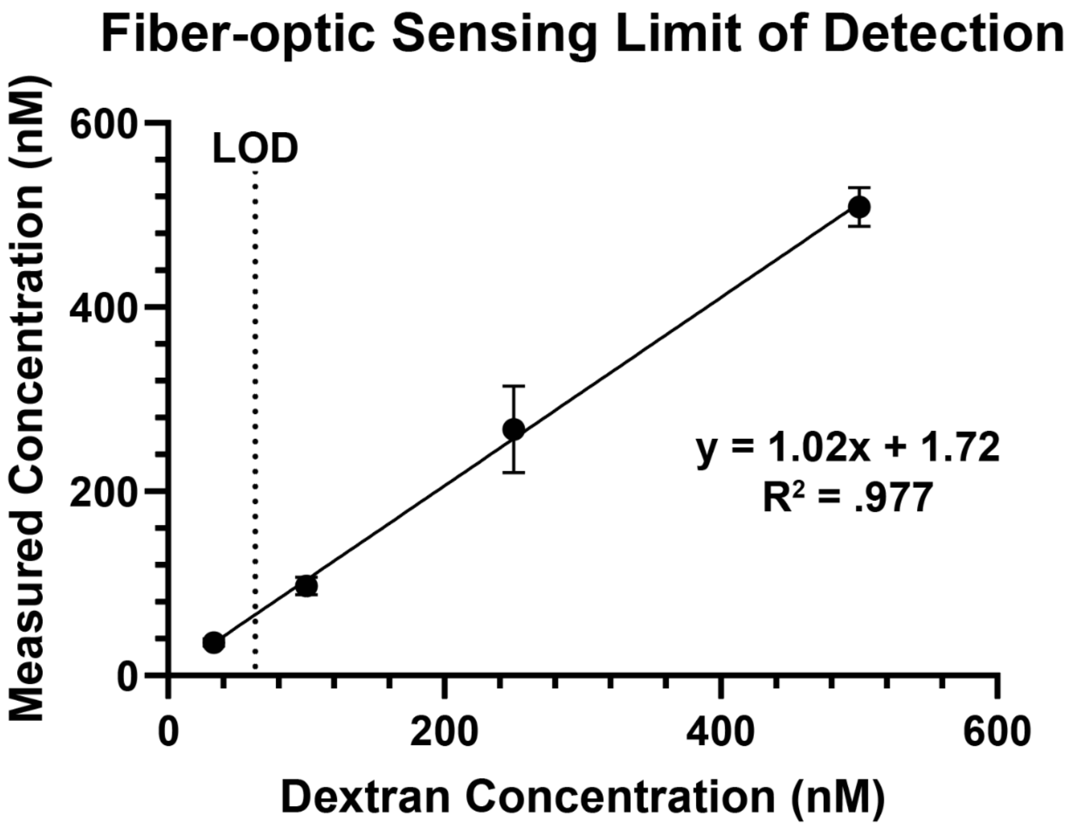


Supplemental Figure 2: Calibration curve obtained using our fiber-optic sensing platform to detect fluorescent dextran of different concentrations on-chip. The measured concentration closely matched the known dextran concentration dosed on-chip. The limit of detection, marked by a horizontal dotted line and labeled with LOD, was determined to be 63 nM.

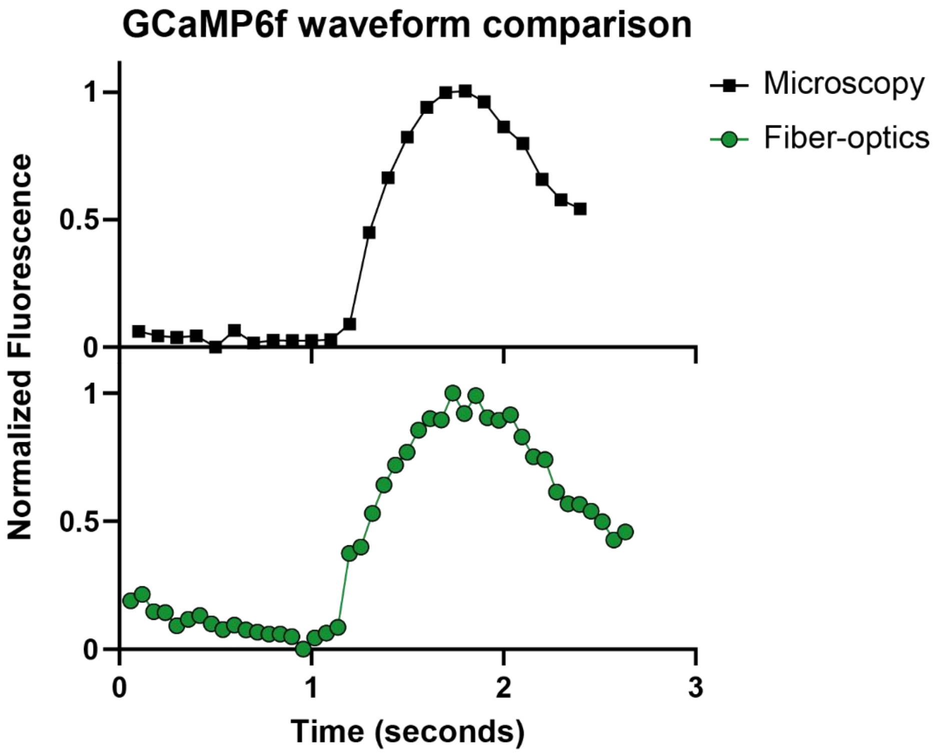


Supplemental Figure 3: Direct GCaMP6f waveform comparison between emission collected by fluorescence microscopy (top) and by our fiber-optic-based sensing platform (bottom). The overall waveform shapes are nearly identical, capturing emitted fluorescence from the exact same cell population of iPSC-derived cardiomyocytes seeded on-chip.

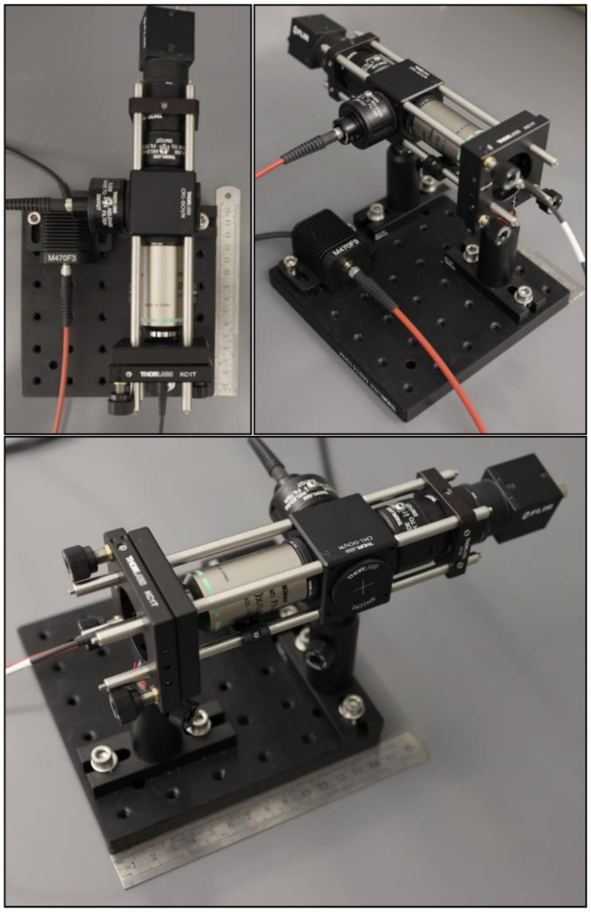


Supplemental Figure 4: Top-down (top left) and trimetric (top right, bottom) photographs of our fiber-optic based sensing platform. Compact optical equipment minimizes total equipment size, with the entire system fitting within a 15 cm x 25 cm x 20 cm volume. Bundled fiber optic cables mounted in the 5-axis translator were routed into an incubator for in situ recording.

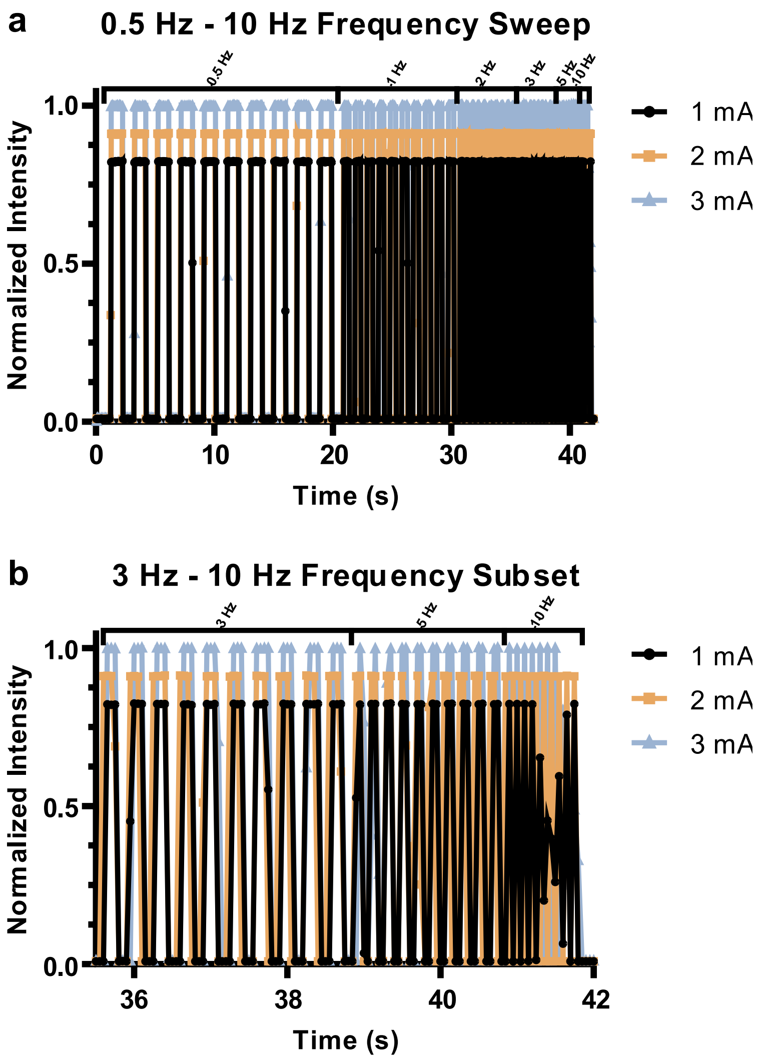


Supplemental Figure 5: a) Resulting signal from capturing LED pulses at 20 HZ using our fiber-optic sensing platform. The CMOS recording camera was locked to 20 Hz recording frequency while a microcontroller sent square-wave TTL triggers to an LED to achieve a 50% duty cycle across a 0.5 – 10 Hz frequency range. 10 pulses were sent at each frequency. b) Subplot of data shown in a) to highlight the impact of higher frequency on recording fidelity. As expected, peaks captured at the Nyquist frequency (10 Hz) do not fully reconstruct the expected waveform.

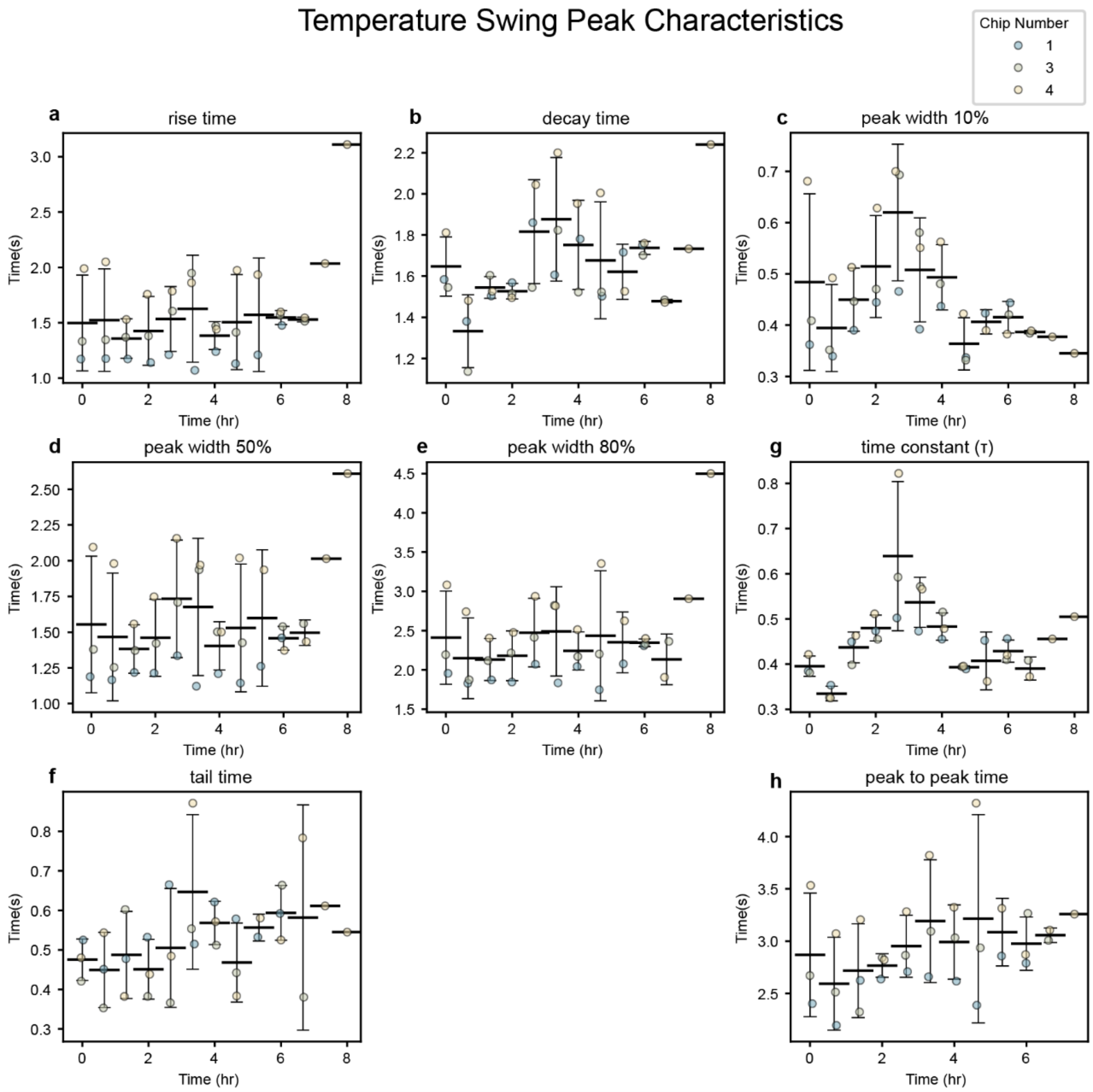


Supplemental Figure 6: a-h) GCaMP6f peak characteristics extracted from each scan during the temperature swing. Each data point represents the average of all active fibers in each chip. Active fiber was defined as a fiber measuring >10 BPM at the first scan (t = 0 hr). Error bars represent mean ± SD, n=12 wells, m = 3 experimental replicates

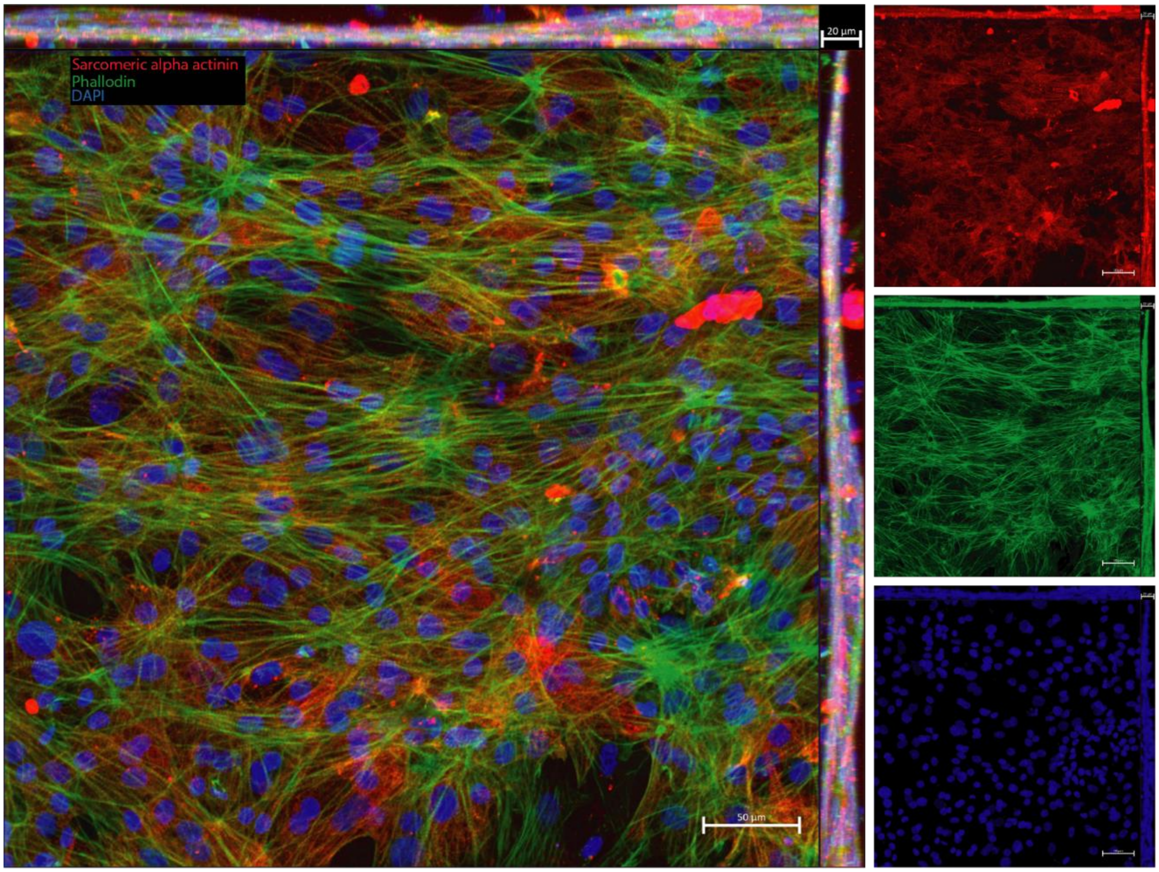


Supplemental Figure 7: Immunofluorescent max projection of hiPSC-derived cardiomyocytes cultured in a single well chip. Cardiomyocytes were cultured for 14 days before being fixed and stained (sarcomeric alpha actinin, red; F-actin, green; nuclei, blue)

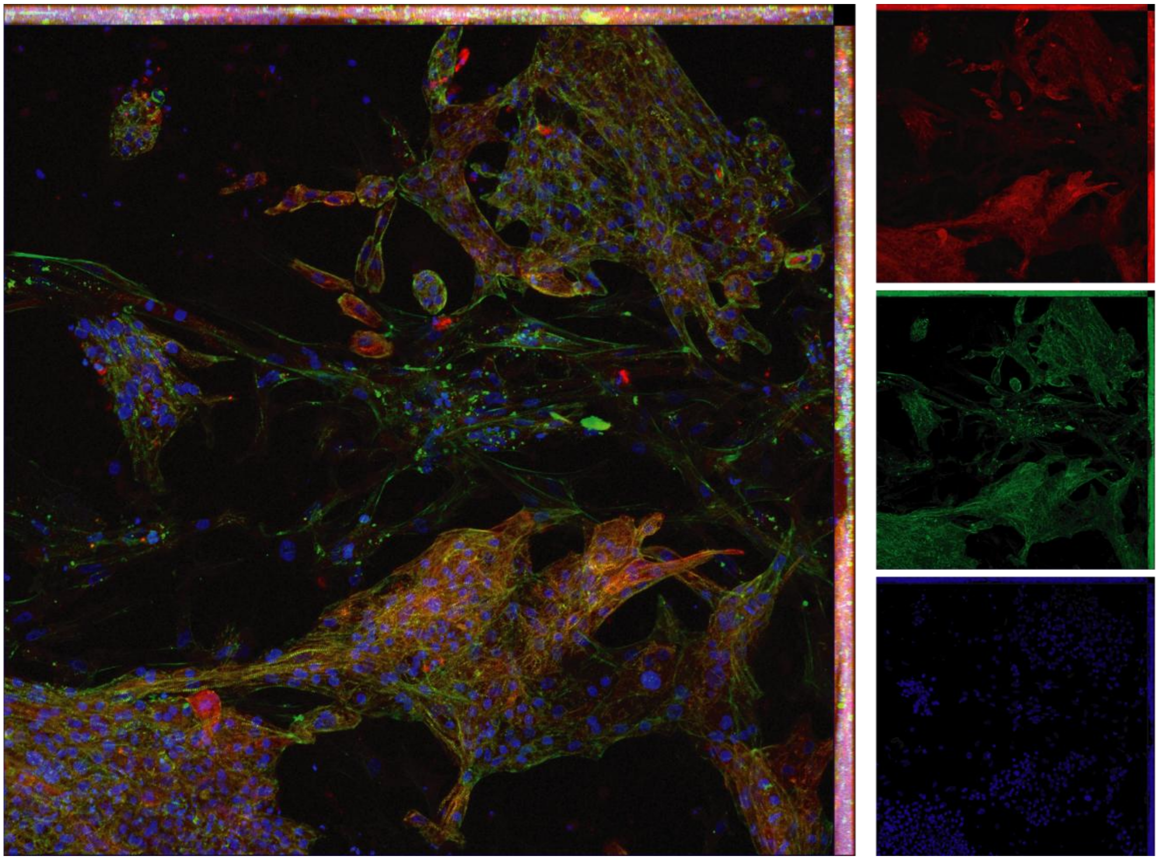

Supplemental Figure 8: Immunofluorescent max projection of hiPSC-derived cardiomyocytes cultured in a single well chip. Cardiomyocytes were cultured for 5 months and 23 days before being fixed and stained (sarcomeric alpha actinin, red; F-actin, green; nuclei, blue)
